# iGS: A Zero-Code Dual-Engine Graphical Software for Polygenic Trait Prediction

**DOI:** 10.64898/2026.02.28.708730

**Authors:** Jiahao Zhang, Fei Chen

**Author notes:** corresponding author Fei Chen.

## Abstract

Genomic selection (GS) has become the core driving force in modern plant and animal breeding. However, state-of-the-art comprehensive GS tools often rely on complex underlying environment configurations and command-line operations, posing significant technical barriers for breeders lacking programming expertise. To address this critical pain point, this study developed a fully “zero-code” graphical user interface (GUI) decision support system for genomic selection. The platform innovatively employs a “portable dual-engine architecture” (R-Portable and Python-Portable) to achieve completely dependency-free, “out-of-the-box” deployment, and integrates a standardized six-step end-to-end workflow from data quality control to result export. Furthermore, the platform comprehensively integrates 33 cutting-edge prediction models across four major paradigms, linear, Bayesian, machine learning, and deep learning, and features an original intelligent parameter configuration system that dynamically renders algorithm parameters to provide a minimalist UI interaction experience. Benchmark testing on the Wheat2000 dataset across six complex agronomic and quality traits, including thousand-kernel weight (TKW) and grain protein content (PROT), demonstrated that classic linear models remain highly robust for polygenic additive traits, while tree-based machine learning and hybrid deep learning architectures exhibit superior predictive potential and noise resilience when resolving complex epistatic effects and low-heritability traits. The successful deployment of this platform fundamentally liberates biologists from the constraints of computational science, providing robust digital infrastructure to accelerate the popularization and practical application of GS technologies in agricultural production.

## Introduction

In modern agricultural breeding systems, genomic selection (GS) has profoundly transitioned from a frontier theoretical exploration into a core operational driver for accelerating the genetic improvement of plants and animals ^1,2^. By utilizing high-density single nucleotide polymorphism (SNP) markers spanning the entire genome, GS technology can directly capture the additive and non-additive genetic variances of all minor-effect polygenes within a target population. This enables the precise estimation of genomic estimated breeding values (GEBVs) without the prerequisite of time-consuming and costly field phenotypic evaluations. This paradigm shift has dramatically shortened the breeding generation interval and significantly increased the genetic gain per unit of time. With the popularization of high-throughput sequencing technologies and the precipitous drop in costs, the primary challenge in the GS domain has shifted from acquiring genotypic data to deciphering the true mapping relationships between complex agronomic traits and massive, high-dimensional genomic data characterized by severe multicollinearity.

To address this challenge, prediction algorithms in computational genetics have undergone rapid iteration and expansion. Early GS research primarily relied on classic linear mixed models (*e*.*g*. GBLUP) and Bayesian inference methods based on various prior assumptions (*e*.*g*. BayesA, BayesB) ^3,4^. In recent years, the introduction of machine learning (ML) and deep learning (DL) architectures has provided entirely new mathematical tools for dissecting complex genomic architectures ^5,6^, such as epistatic effects and genotype-by-environment (GxE) interactions. Against the backdrop of this algorithmic proliferation, researchers have sought to integrate diverse, heterogeneous models into unified analytical frameworks. Currently, the most representative competitive product in this domain is MultiGS ^7^. The MultiGS framework constructs two complementary computational pipelines, MultiGS-R (based on Java/R) and MultiGS-P (based on Python), successfully establishing a standardized workflow for model evaluation.

However, despite the significant achievements of tools like MultiGS in algorithm integration, their widespread adoption in practical breeding applications is severely hindered by “software engineering and dependency barriers”. The execution of MultiGS heavily relies on intricate underlying environment configurations. Users are required to manually configure Java runtime environments, independently install R interpreters, and compile necessary dependency packages via the command line. For deep learning modules ^8,9^, users must possess specialized skills to configure Python virtual environments and manage PyTorch dependencies under Linux ecosystems. For front-line breeders who generally lack support from dedicated bioinformatics teams, such code-dependent command-line interface (CLI) tools constitute an exceedingly high technical threshold.

To thoroughly eliminate the dependency pain points associated with deploying computational tools in practical breeding, this study developed and validated a novel, completely zero-code GUI genomic selection platform ^10,11^. Distinguishing itself fundamentally from MultiGS, this platform leverages an innovative “Dual-Engine Architecture” to encapsulate all R packages, Python scientific computing libraries, and their corresponding environmental dependencies into fully portable modules. These are seamlessly migrated onto a highly interactive GUI. Users can invoke up to 33 state-of-the-art prediction models within a unified decision support system without writing a single line of code or installing third-party environments ^12,13^.

## Materials and Methods

### Software Engineering and System Development

The front-end graphical interactive interface of the platform was constructed using the Python PyQt5 framework, utilizing the Signal and Slot mechanism to achieve asynchronous, non-blocking communication between the front and back ends. To realize a fully “zero-code and dependency-free” deployment, a dynamic path resolution and execution state detection mechanism (get_base_paths) was developed at the system’s core. The system internally integrates a portable R environment (R-Portable) and a portable Python environment (Python-Portable). During the execution of computational tasks, the GUI core scheduler employs the subprocess module to dispatch standardized parameter protocols directly to isolated sandbox engines within the resource directory (RES_DIR), thereby circumventing any contamination of or reliance on the host operating system’s environment variables.

### Evaluation Dataset (Wheat2000)

To objectively assess the predictive performance of the 33 built-in models, this study utilized the internationally recognized Wheat2000 dataset ^14,15^ as the benchmark material^16,17^. This dataset comprises phenotypic and genotypic data for 2,000 elite bread wheat landraces. During the genotype quality control phase, the system invoked the underlying PLINK engine, setting the minor allele frequency (MAF) threshold to 0.05 and the maximum missing rate to 0.2, ultimately retaining 9,927 high-quality SNP markers for downstream analysis ^18^. The study evaluated six representative agronomic and quality traits characterized by diverse genetic architectures: thousand-kernel weight (TKW), test weight (TW), grain width (WIDTH), grain length (LENGTH), grain hardness (HARD), and grain protein content (PROT).

### Cross-Validation and Model Evaluation Strategy

To ensure the rigor and reproducibility of the benchmark testing, the Wheat2000 population was strictly partitioned into a training set of 1,600 samples and an independent test set of 400 samples based on a standard 80:20 ratio using a fixed random seed. The predictive accuracy of the models was quantified by calculating the Pearson Correlation Coefficient (PCC) between the predicted GEBVs and the true phenotypic observations in the test set ^19^. During the benchmark evaluation, a standardized hyperparameter tuning process using Grid Search was conducted across all models to ensure a fair comparison.

## Results

### System Architecture and Algorithm Implementation

Unlike MultiGS, which fragments data preprocessing, model training, and result evaluation into separate scripts, this platform implements a highly standardized “end-to-end” decision support workflow directly at the GUI level. This process is meticulously designed as six sequential, standardized steps covering the entire lifecycle from raw sequencing data to final breeding value outputs.

As illustrated in Figure 1, the system’s input layer supports VCF files, phenotypic data tables, and preprocessed 0/1/2 matrices. Upon entering the integrated GUI core, the system guides the user from left to right through the following steps: Quality Control (QC): Invokes the underlying PLINK engine to execute MAF and missing rate filtering; Genotype Imputation: Performs missing value imputation on the filtered genotype matrix; Population Structure Analysis: Executes Principal Component Analysis (PCA) and renders 2D/3D structural clustering plots; GWAS: Estimates marker effects and generates Manhattan and QQ plots; Genomic Prediction Engine: The core computational hub of the system. Leveraging the R+Python dual-engine architecture, it concurrently schedules linear models, machine learning, deep learning, and hybrid architectures; Result Integration & Export: Automatically generates prediction accuracy scatter plots, loss curves, and GEBV tables, supporting one-click exportation.

**Figure 1.**
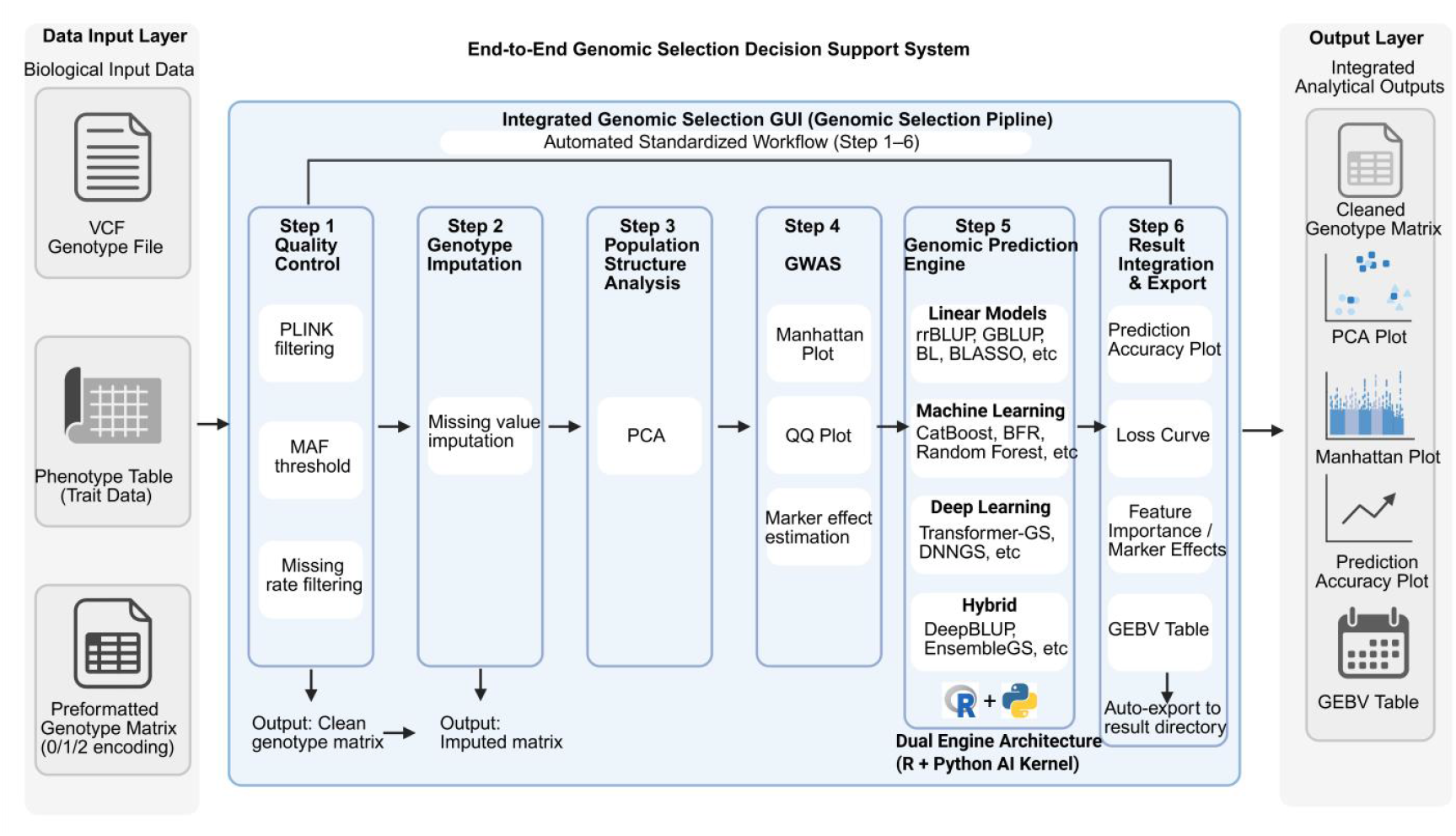
Architecture of the End-to-End Genomic Selection Decision Support System and the standardized six-step workflow. The left panel displays the Data Input Layer, supporting raw VCF files, phenotypic tables, and genotype matrices. The central core area illustrates the automated GUI pipeline comprising: (1) Quality Control (PLINK-based MAF and missing rate filtering), (2) Genotype Imputation, (3) Population Structure Analysis (PCA), (4) GWAS (Manhattan and QQ plots), (5) Genomic Prediction Engine (the core R+Python Dual-Engine architecture orchestrating various models), and (6) Result Integration & Export. The right panel represents the Output Layer, showcasing automatically generated comprehensive analytical charts and GEBV reports.

### Classification and Mathematical Features of the 33 Prediction Models

To address traits ranging from simple oligogenic structures to highly environmentally sensitive polygenic networks, this platform globally integrates an unprecedented 33 genomic prediction models into a unified UI system. These models are rigorously categorized into four major technical paradigms, as detailed in Table 1.

**Table 1.**
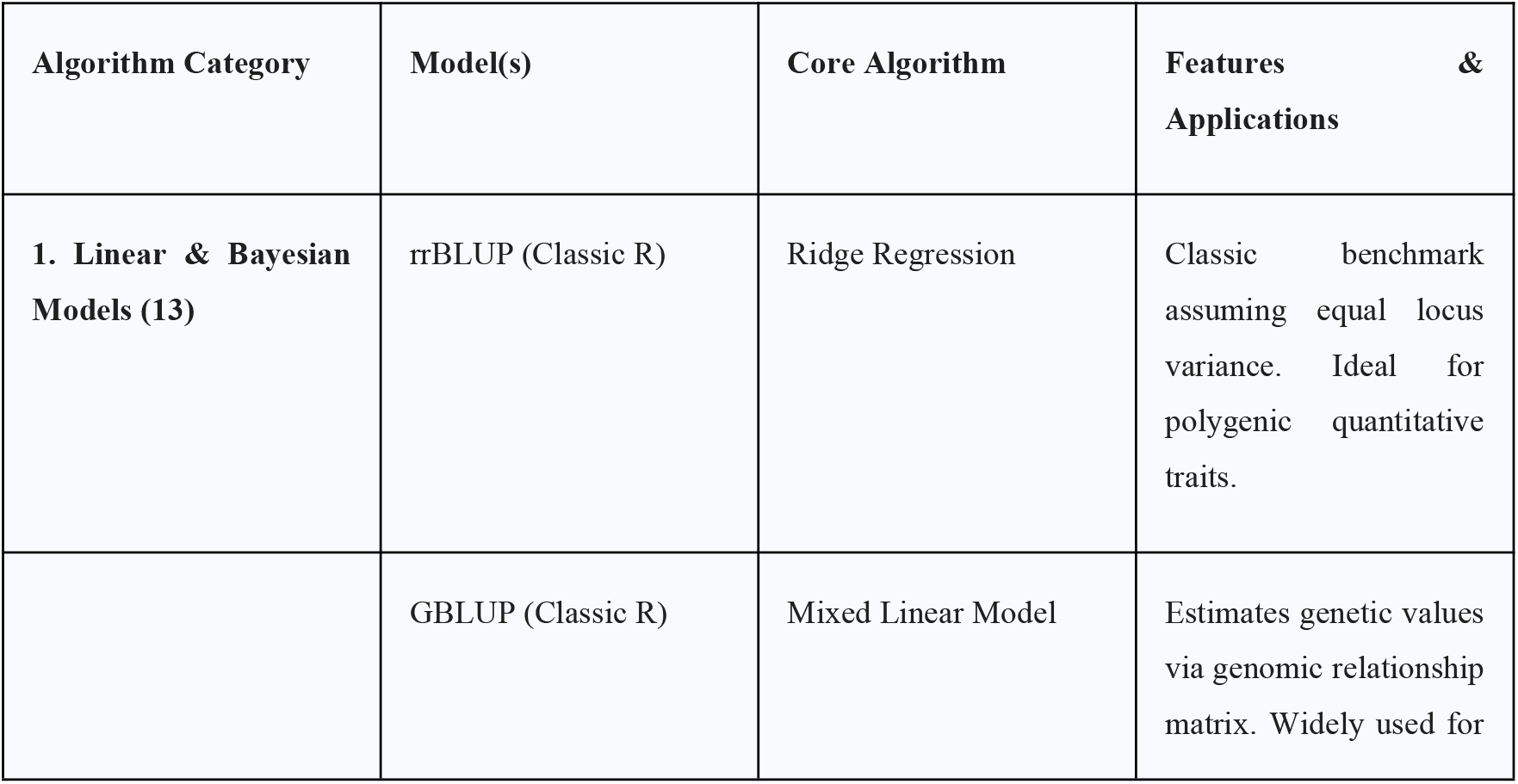
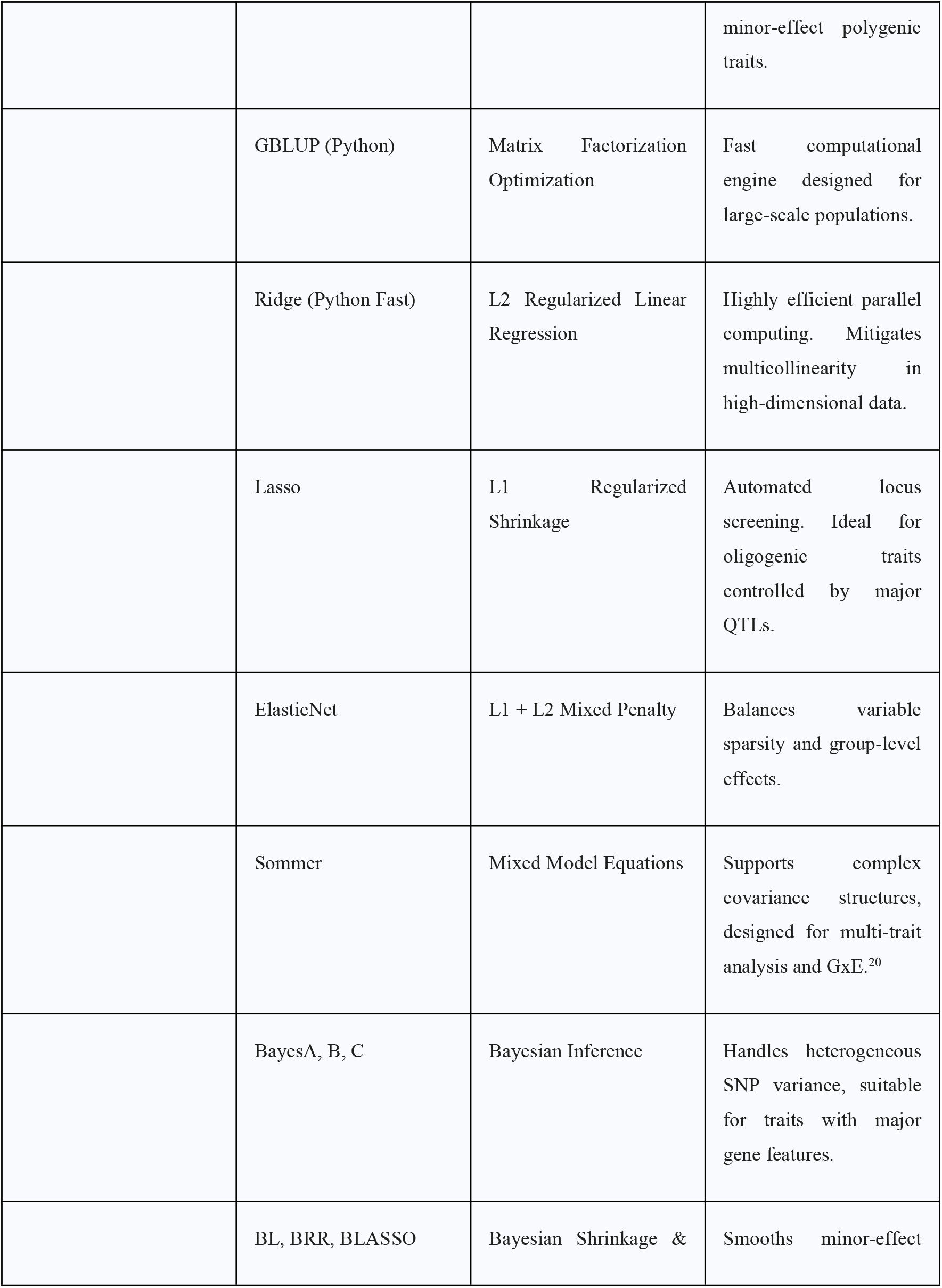
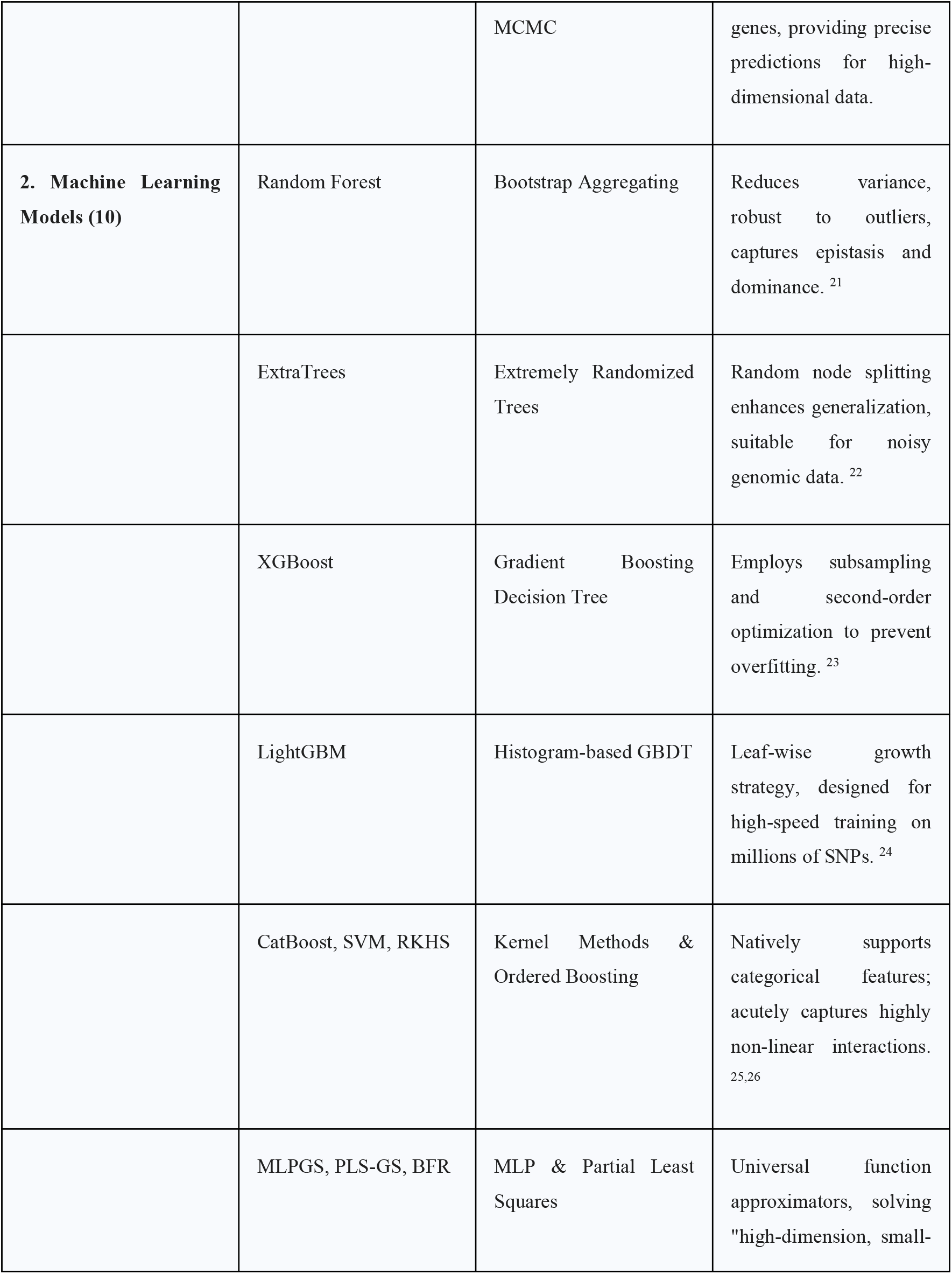
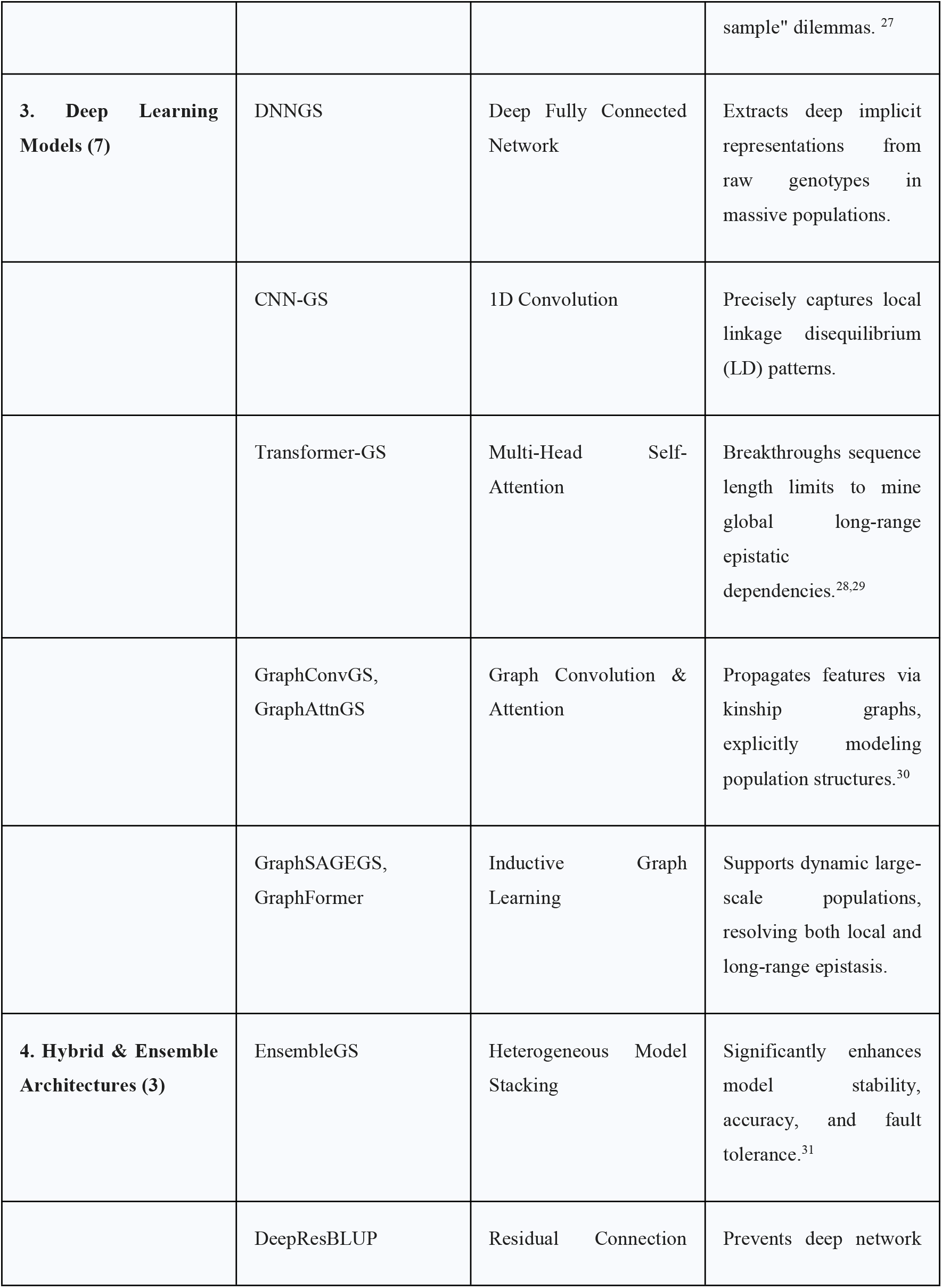
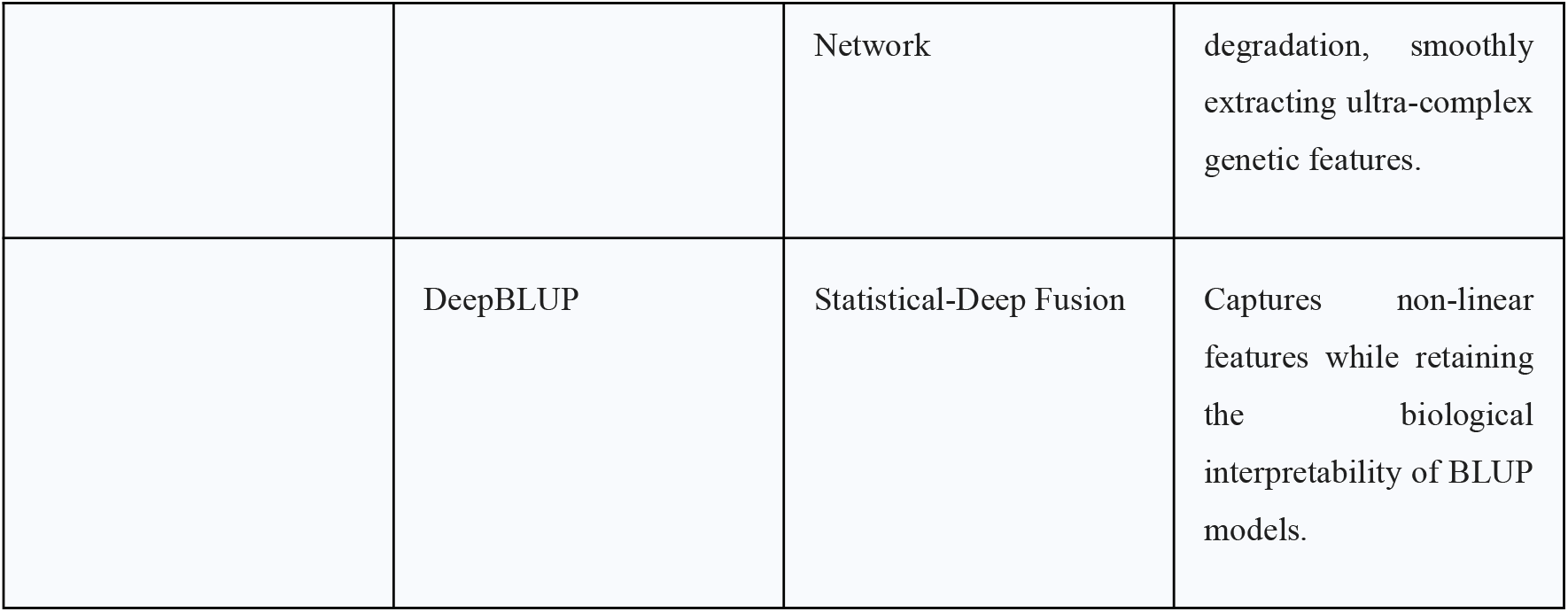
Summary of the 33 genomic prediction models integrated into the platform.

### Intelligent Parameter Protocol and Dynamic Configuration System

Exposing the hyper-parameters of 33 mathematically disparate models simultaneously would trigger cognitive overload for users. Consequently, the platform introduces an innovative “Intelligent Parameter Configuration System” based on the GUI (Figure 2).

**Figure 2.**
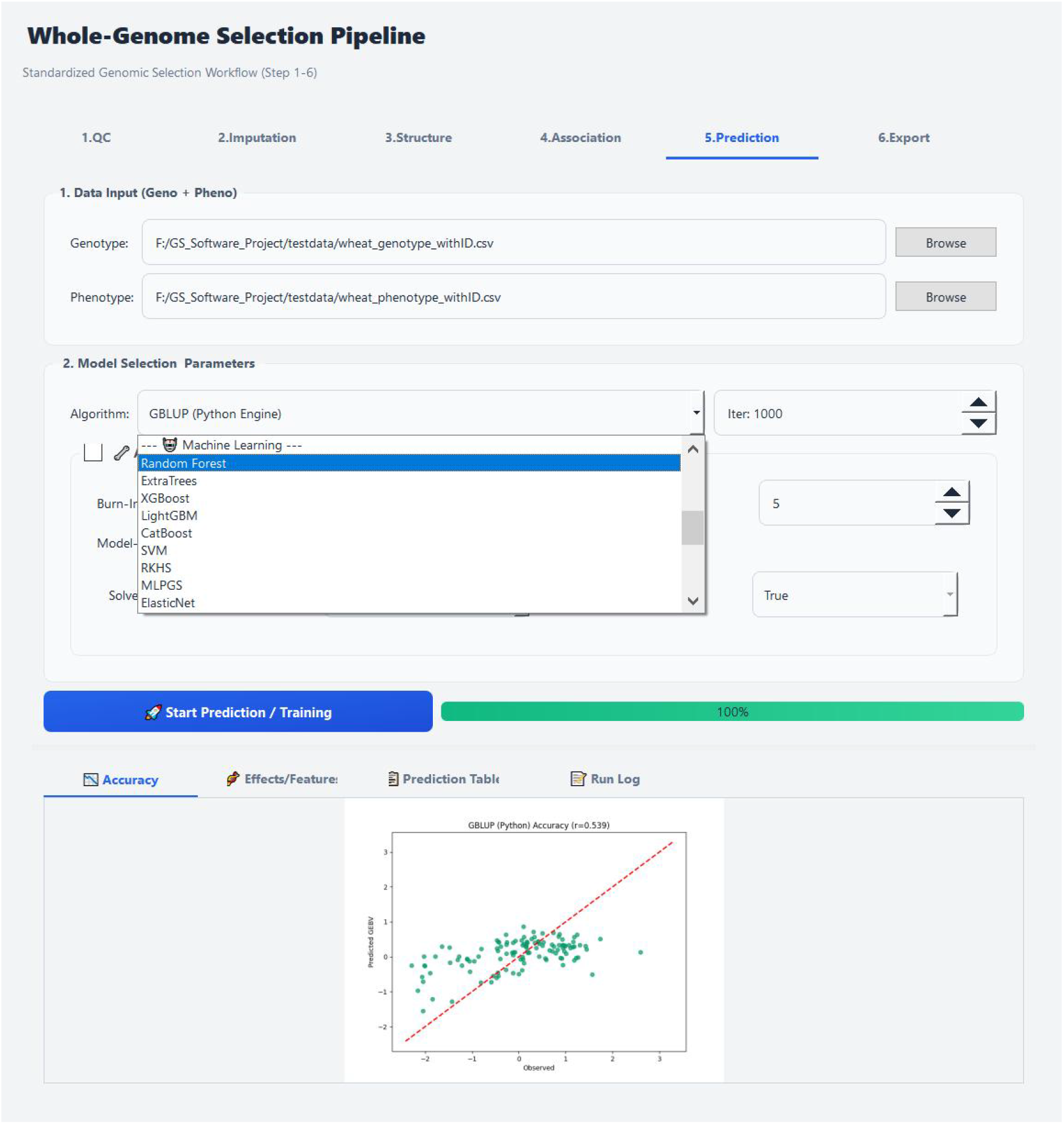
Graphical User Interface (GUI) of the whole-genome selection platform featuring the intelligent dynamic parameter configuration system. The screenshot highlights “Step 5: Prediction” of the standardized workflow. The upper section provides file input interfaces and an intuitive algorithm selection drop-down menu (currently displaying the Machine Learning models list). The right section features a “model-aware” dynamic parameter panel, where the system automatically renders core hyperparameters matching the selected model (*e*.*g*. Iterations) while hiding irrelevant ones. The lower section displays a real-time interactive results panel, showcasing the prediction accuracy scatter plot upon training completion, fully embodying the “zero-code, dependency-free” minimalist user experience.

Regardless of the model selected by the front-end user, the GUI layer serializes the complex configuration panel into a standardized 6-element parameter string protocol: “Iter|LR|Depth|Batch|Dropout|BurnIn”. This is subsequently transmitted via underlying worker scripts to the isolated AI engines.

As shown in Figure 2, during the genomic prediction phase (Step 5), users can freely switch between different algorithmic paradigms (e.g., Random Forest under Machine Learning) using an intuitive drop-down menu (Algorithm). The system’s back-end implements “Model-Aware” dynamic visibility control: upon selecting a specific model, the right-side interface automatically renders matched core parameters (e.g., Iter: 1000) while hiding irrelevant ones (e.g., hiding Bayesian-exclusive Burn-In parameters when XGBoost is selected). Post-training, the system seamlessly renders prediction accuracy scatter plots (e.g., GBLUP Python engine Accuracy chart) within the same interface, achieving a genuinely zero-code interactive workflow.

### Benchmark Test Results and Analysis

#### Multi-Trait Prediction Performance Evaluation on the Wheat2000 Dataset

To visually demonstrate the performance of various models under the dual-engine architecture, a systematic evaluation of prediction performance was conducted on six complex agronomic and quality traits using the Wheat2000 dataset. As depicted in Figure 3, the x-axis represents prediction accuracy (Pearson Correlation Coefficient), and the y-axis illustrates the distribution of the 29 models across the four major categories (Linear, ML, DL, Hybrid). Four Graph Neural Network (GNN) models were excluded from this benchmark because standard PCA dimensionality reduction disrupts the topological dependencies required for graph construction.

**Figure 3.**
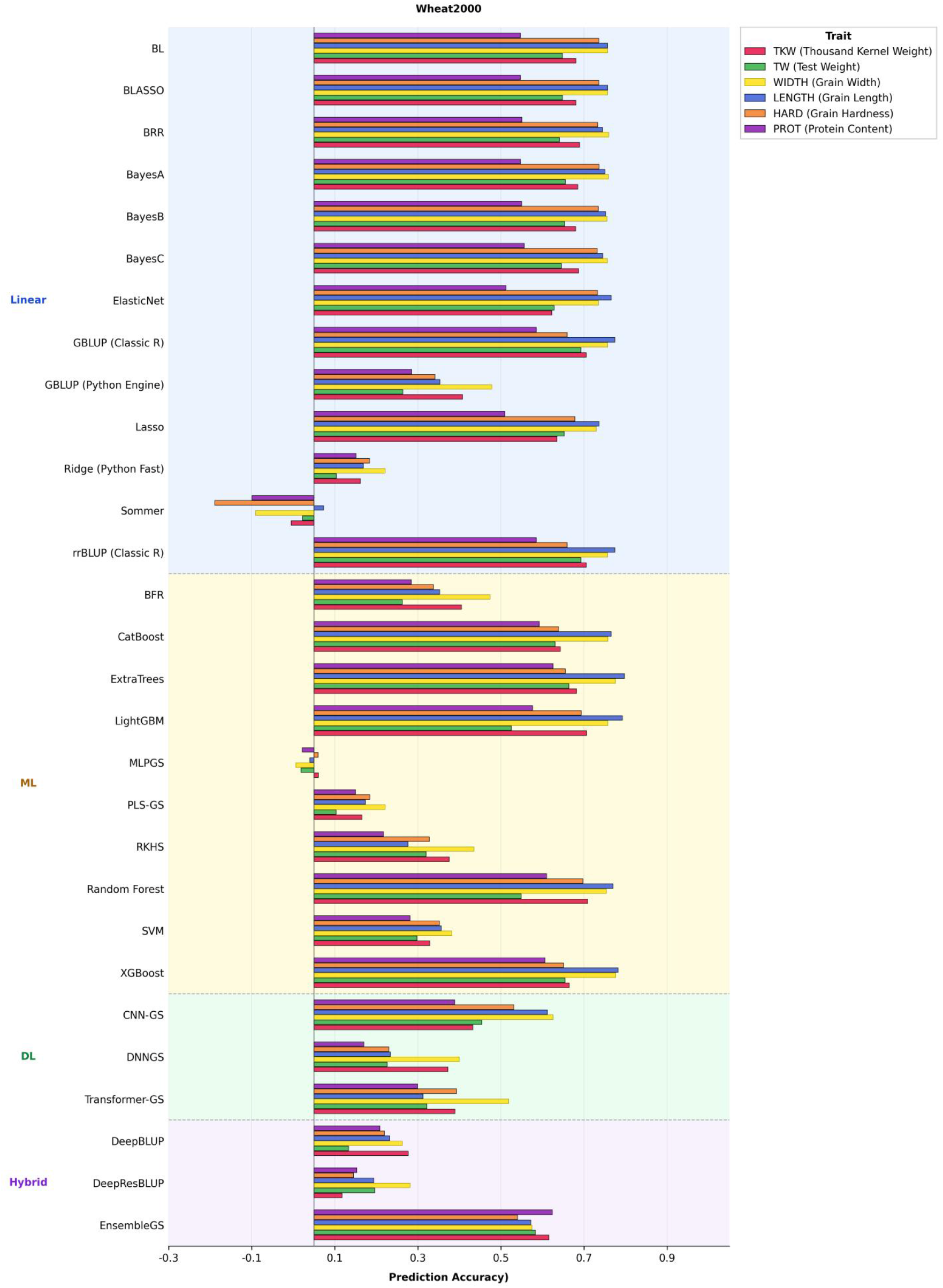
Evaluation of prediction accuracy (Pearson Correlation Coefficient) across six agronomic and quality traits using the Wheat2000 dataset. The bar chart illustrates the performance of 29 models (Four graph-topology-based models were excluded from this benchmark as the preceding PCA dimensionality reduction flattens the data, destroying the spatial topological structures required for their inputs.) distributed across four major algorithmic paradigms: Linear/Bayesian, Machine Learning (ML), Deep Learning (DL), and Hybrid architectures. The evaluated traits include thousand-kernel weight (TKW), test weight (TW), grain width (WIDTH), grain length (LENGTH), grain hardness (HARD), and protein content (PROT). Theoretical note on untested models: Four Graph Neural Network (GNN) models (GraphConvGS, GraphAttnGS, GraphSAGEGS, GraphFormer) integrated within the system were excluded from this benchmark. Theoretically, while traditional ML/DL models process flattened 2D matrices (0/1/2 SNP arrays or PCA-reduced features), GNNs operate on non-Euclidean data. They require inputs to be reconstructed into complex spatial topological graphs comprising nodes (samples) and edges (kinship/LD interactions). The standard PCA dimensionality reduction destroys the topological dependencies required for graph construction. Practically, computing these topological graphs relies on the torch_geometric library, which requires complex, hardware-specific C++/CUDA compilation. To preserve the absolute portability and zero-configuration compatibility of the system’s dual-engine architecture on standard PCs, this heavy dependency was not bundled in the base environment (triggering a ModuleNotFoundError during automated batching). Consequently, these graph-topology-based models were excluded from this flat-matrix feature benchmark.

#### Prediction Performance for Yield Components and Highly Polygenic Traits

For continuous quantitative traits such as thousand-kernel weight (TKW), test weight (TW), grain width (WIDTH), and grain length (LENGTH), which are typically controlled by the cumulative superposition of numerous minor-effect genes, the classic Linear Models paradigm demonstrated its robustness as the gold standard. rrBLUP and its Python-accelerated equivalent (Ridge Python Fast) consistently maintained high prediction accuracies between 0.70 and 0.78 for traits like WIDTH and LENGTH. This proves that in genetic architectures dominated by pure additive variance, the L2 regularization penalty mechanism perfectly aligns with the underlying biological logic.

However, tree-ensemble algorithms within the Machine Learning paradigm further breached the ceiling of linear models. In predicting TKW and LENGTH, the accuracies of ExtraTrees, XGBoost, and LightGBM generally exceeded 0.75, with some approaching 0.80. This highlights the existence of non-negligible non-additive effects (*e*.*g*. epistatic interactions between genes) underlying yield components, and gradient boosting decision trees can implicitly reconstruct these complex interactive networks in a non-parametric manner.

#### Prediction Performance for Complex Quality and Low-Heritability Traits (HARD, PROT)

Compared to yield traits, grain hardness (HARD) and protein content (PROT) are often subject to intense environmental interactions and possess relatively lower heritability. As clearly observed in Figure 3, the accuracies of all models for HARD and PROT generally regressed to the 0.4 to 0.65 range.

In such highly challenging scenarios, the Hybrid Architecture and Bayesian paradigm demonstrated unique noise-resistance advantages. EnsembleGS, which employs heterogeneous model stacking, firmly occupied the top tier in predicting HARD and PROT, substantiating the engineering value of model ensembling when dealing with high-noise traits. Simultaneously, Bayesian models with strong variable selection capabilities, such as BayesB, outperformed standard GBLUP by applying shrinkage to ineffective background SNPs. While certain Deep Learning models (*e*.*g*. Transformer-GS30) performed remarkably well on traits like TW and WIDTH, simple multi-layer perceptrons (*e*.*g*. MLPGS without residual connections) were prone to overfitting under small-sample constraints, as evidenced by MLPGS’s bottom-tier performance on specific traits in Figure 3. This underscores the critical importance of integrating diverse architectures within the platform to allow breeders to conduct cross-validation.

## Discussion and Conclusion

The deep popularization of genomic selection technology is the core engine driving the transition of modern agriculture from “empirical breeding” to “precision design breeding”^32,33^. Historically, outstanding academic tools like MultiGS, despite proving the potential of DL hybrid architectures in cross-population predictions, imposed deployment barriers requiring users to navigate the disparate ecosystems of Java, R, and Python, severely impeding the technology’s integration into the industry.

The zero-code dual-engine platform developed in this study thoroughly dismantles these barriers through its innovative portable environment mounting mechanism (R-Portable/Python-Portable) and intelligent UI parameter protocols. The benchmark testing on the Wheat2000 dataset robustly refutes the illusion of a “universal optimal model”: linear models remain robust for additive traits, tree models excel at capturing non-additive variances, and hybrid deep architectures show great potential in complex noise-resilient tasks. Through a highly integrated GUI, breeders can now bypass tedious code debugging, invoking 33 top-tier algorithms with a single click, thereby refocusing their invaluable time and energy on elucidating the biological mechanisms of traits and formulating breeding strategies. The inception of this platform marks the official entry of genomic selection tools into an era of zero-code, dependency-free popularization.

## Author Contributions

F.C. designed this research. Y.Z. participated in the design of this project and the writing of the draft manuscript. J.Z. performed the analyses and wrote the draft manuscript. All authors read and approved the final manuscript.

## Acknowledgments

We sincerely acknowledge the support of the FAN system provided by the Precision Design and Intelligent Manufacturing Platform of Yazhouwan National Laboratory, which greatly facilitated this work.

## Data and Code Availability

Datasets used in this research were from publicly available sources as described in Materials and Methods. The software and various utility programs for this research are freely available on GitHub (https://github.com/waterlilyfeichen-afk/iGS-Breeding).

## References

1 Meuwissen, T. H., Hayes, B. J. & Goddard, M. Prediction of total genetic value using genome-wide dense marker maps. genetics 157, 1819–1829 (2001).

2 Wallace, J. G., Rodgers-Melnick, E. & Buckler, E. S. On the road to breeding 4.0: unraveling the good, the bad, and the boring of crop quantitative genomics. Annual review of genetics 52, 421–444 (2018).

3 VanRaden, P. M. Efficient methods to compute genomic predictions. Journal of dairy science 91, 4414–4423 (2008).

4 Gianola, D. Priors in whole-genome regression: the Bayesian alphabet returns. Genetics 194, 573–596 (2013).

5 Xu, Y. et al. Smart breeding driven by big data, artificial intelligence, and integrated genomic-enviromic prediction. Molecular plant 15, 1664–1695 (2022).

6 Sandhu, K. S., Lozada, D. N., Zhang, Z., Pumphrey, M. O. & Carter, A. H. Deep learning for predicting complex traits in spring wheat breeding program. Frontiers in plant science 11, 613325 (2021).

7 You, F. M. et al. MultiGS: A comprehensive and user-friendly genomic prediction platform Integrating statistical, machine learning, and deep learning models for breeders. bioRxiv, 2026.2001.2002.697306 (2026).

8 Pérez-Enciso, M. & Zingaretti, L. M. A guide on deep learning for complex trait genomic prediction. Genes 10, 553 (2019).

9 Ma, W. et al. A deep convolutional neural network approach for predicting phenotypes from genotypes. Planta 248, 1307–1318 (2018).

10 Mu, H. et al. OmicShare tools: a zero-code interactive online platform for biological data analysis and visualization. Imeta 3, e228 (2024).

11 Jubair, S. & Domaratzki, M. in 2019 IEEE International Conference on Bioinformatics and Biomedicine (BIBM). 1993-2000 (IEEE).

12 Endelman, J. B. Ridge regression and other kernels for genomic selection with R package rrBLUP. The plant genome 4 (2011).

13 Pérez, P. & de Los Campos, G. Genome-wide regression and prediction with the BGLR statistical package. Genetics 198, 483–495 (2014).

14 González, F., García-Abadillo, J. & Jarquín, D. Introducing CHiDO—A No Code Genomic Prediction software implementation for the characterization and integration of driven omics. The Plant Genome 18, e20519 (2025).

15 Chen, C. et al. PlantMDCS: A code-free, modular toolkit for rapid deployment of plant multiomics databases. bioRxiv, 2026.2002.2009.704752 (2026).

16 Chao, D. et al. Mtcro: multi-task deep learning framework improves multi-trait genomic prediction of crops. Plant Methods 21, 12 (2025).

17 Liu, Y. et al. Phenotype prediction and genome-wide association study using deep convolutional neural network of soybean. Frontiers in genetics 10, 1091 (2019).

18 Amadeu, R. R. et al. AGHmatrix: R package to construct relationship matrices for autotetraploid and diploid species: a blueberry example. The plant genome 9, plantgenome2016.2001.0009 (2016).

19 Li, J. et al. TrG2P: A transfer-learning-based tool integrating multi-trait data for accurate prediction of crop yield. Plant Communications 5 (2024).

20 Covarrubias-Pazaran, G. Genome-assisted prediction of quantitative traits using the R package sommer. PloS one 11, e0156744 (2016).

21 Breiman, L. Random forests. Machine learning 45, 5–32 (2001).

22 Geurts, P., Ernst, D. & Wehenkel, L. Extremely randomized trees. Machine learning 63, 3–42 (2006).

23 Chen, T. & Guestrin, C. in Proceedings of the 22nd acm sigkdd international conference on knowledge discovery and data mining. 785–794.

24 Ahmad, N., Wali, B. & Khattak, A. J. Heterogeneous ensemble learning for enhanced crash forecasts–A frequentist and machine learning based stacking framework. Journal of safety research 84, 418–434 (2023).

25 Prokhorenkova, L., Gusev, G., Vorobev, A., Dorogush, A. V. & Gulin, A. CatBoost: unbiased boosting with categorical features. Advances in neural information processing systems 31 (2018).

26 Gianola, D., Fernando, R. L. & Stella, A. Genomic-assisted prediction of genetic value with semiparametric procedures. Genetics 173, 1761–1776 (2006).

27 Rigdon, E. E. Rethinking partial least squares path modeling: In praise of simple methods. Long range planning 45, 341–358 (2012).

28 Vaswani, A. et al. in Neural Inf. Process. Syst. 5999–6009.

29 Wu, C. et al. A transformer-based genomic prediction method fused with knowledge-guided module. Briefings in Bioinformatics 25, bbad438 (2024).

30 Kipf, T. N. & Welling, M. Semi-supervised classification with graph convolutional networks. arXiv preprint arXiv:1609.02907 (2016).

31 Tomura, S., Wilkinson, M. J., Cooper, M. & Powell, O. Improved genomic prediction performance with ensembles of diverse models. G3: Genes, Genomes, Genetics 15, jkaf048 (2025).

32 Zhu, W. et al. The CropGPT project: Call for a global, coordinated effort in precision design breeding driven by AI using biological big data. Molecular Plant 17, 215–218 (2024).

33 Liu, J., Fernie, A. R. & Yan, J. Crop breeding–From experience-based selection to precision design. Journal of plant physiology 256, 153313 (2021).

